# Polo-like kinase 4 (Plk4) potentiates *anoikis*-resistance of p53KO mammary epithelial cells by inducing a hybrid EMT phenotype

**DOI:** 10.1101/2022.12.16.520613

**Authors:** Irina Fonseca, Cíntia Horta, Ana Sofia Ribeiro, Barbara Sousa, Gaëlle Marteil, Mónica Bettencourt-Dias, Joana Paredes

## Abstract

Polo-like kinase 4 (Plk4), the major regulator of centriole biogenesis, has emerged as a putative therapeutic target in cancer due to its abnormal expression in human carcinomas, leading to centrosome number deregulation, mitotic defects and chromosomal instability. Moreover, Plk4 deregulation promotes tumor growth and metastasis in mouse models and is significantly associated with poor patient prognosis.

Here, we further investigate the role of Plk4 in carcinogenesis and show that its overexpression significantly potentiates resistance to cell death by *anoikis* of non-tumorigenic p53 knock-out (p53KO) mammary epithelial cells. Importantly, this effect is independent of Plk4’s role in centrosome biogenesis, suggesting that this kinase has additional cellular functions. Interestingly, the Plk4-induced *anoikis* resistance is associated with the induction of a stable hybrid epithelial-mesenchymal phenotype and is partially dependent on P-cadherin upregulation. Furthermore, we found that the conditioned media of Plk4-induced p53KO mammary epithelial cells also induces *anoikis* resistance of breast cancer cells in a paracrine way, being also partially dependent on soluble P-cadherin secretion.

Our work shows, for the first time, that high expression levels of Plk4 induce *anoikis* resistance of both mammary epithelial cells with p53KO background, as well as of breast cancer cells exposed to their secretome, which is partially mediated through P-cadherin upregulation. These results reinforce the idea that Plk4, independently of its role in centrosome biogenesis, functions as an oncogene, by impacting the tumor microenvironment to promote malignancy.

## Introduction

Centrosomes are the primary microtubule-organizing center (MTOC) of dividing animal cells. Consisting of two centrioles surrounded by a pericentriolar matrix (PCM), they participate in essential cellular processes, such as cell division, signaling, migration, and polarity(1,2). Centriole biogenesis is tightly regulated and occurs once per cell cycle, in S-phase, during which centrioles duplicate to ensure the assembly of a bipolar spindle in mitosis, an essential structure for proper chromosome segregation(2–5).

Polo-like kinase 4 (Plk4), is a regulator of centriole biogenesis that is required for centriole duplication(5–8) via the phosphorylation and interaction with different centriolar proteins. In consequence, when Plk4 is overexpressed, it leads to centrosome amplification (CA, i.e., the presence of supernumerary centrosomes) resulting in cells with more than 4 centrioles(9), promoting genomic instability and tumorigenesis(10–15). In fact, Plk4 overexpression, as well as CA and centrosome structure abnormalities, have been already found in many types of tumors(14–19) and shown to be significantly associated with poor clinical outcomes for cancer patients(18,20–22).

Despite strong evidence of Plk4’s role in cancer and its therapeutic potential, it is still unclear how it contributes to tumorigenesis. Plk4 has been demonstrated to enhance cell migration through RhoA activation and to be associated with the expression of matrix metalloproteinases (MMPs) and other pro-motility genes(23). In addition, Plk4 has been shown to promote cancer metastasis through the regulation of the actin cytoskeleton by the Arp2/3 complex(24) in breast cancer cells, whereas its downregulation suppresses tumorigenesis and metastasis in neuroblastoma(25). Plk4 has also been shown to have a prognostic value, since its overexpression is associated with worse disease-free survival (DFS) and overall survival (OS), demonstrating also a negative impact on the response to chemotherapy(22,26,27). Levine *et al*. showed that transient Plk4 overexpression causes CA and aneuploidy *in vivo* in a mouse model of intestinal neoplasia and accelerates the occurrence of spontaneous tumor formation(20), and Serçin *et al*. demonstrated that transient Plk4 overexpression accelerates tumorigenesis in p53-deficient mice(28).

The loss of the p53 tumor suppressor gene has been strongly associated with tumorigenesis and malignant progression in the majority of epithelial cancers(29–31). One of the molecular mechanisms potentiated by the absence of p53 is the induction of epithelial to mesenchymal transition (EMT)(32,33). EMT is the process by which cells lose their epithelial characteristics and undergo several molecular, biochemical, and morphological changes, acquiring a more undifferentiated and mesenchymal phenotype. EMT is well known for conferring stem-like properties and cell plasticity, leading to the acquisition of a migratory and invasive phenotype by weakening cell-cell adhesion, facilitating metastatic capacity, as well as resistance to chemo and radiotherapy(34–36). EMT also allows evasion to *anoikis*, a specific mode of apoptotic cell death that occurs due to insufficient cell-matrix interactions(37,38). Resistance to *anoikis* is a characteristic found in cells with stem-like properties, being thus, considered a critical contributor to cancer cells dissemination within the bloodstream and, consequently, to their metastatic capacity(39,40). Interestingly, high Plk4 expression has been shown to induce EMT in neuroblastoma and colorectal cancer by regulating PI3K/Akt and Wnt/ß-catenin signaling pathways, respectively(25,41), which are well known oncogenic pathways.

Although EMT has been initially assumed as a binary process, the concept of hybrid epithelial-mesenchymal phenotype has recently emerged and has been shown to be more relevant than previously thought for cells to become metastatic. By expressing both epithelial and mesenchymal markers, cells increase their plasticity and are able to better respond to external stimuli, potentiating their capacity to resist apoptosis (as *anoikis*) and enhancing their metastatic features(36,42). P-cadherin, which is a cell-cell adhesion protein, is expressed in cancer cells harboring epithelial and mesenchymal features, being a putative marker of a hybrid EMT phenotype(43). In fact, P-cadherin expression is early promoted by EMT-inducers, such as hypoxia, driving *anoikis* resistance capacity in breast cancer cells(44). Moreover, its overexpression has been found in many types of tumors(45–47) and is associated with tumorigenic and metastatic potential, stem cell activity, and collective cell invasion(48–50). For all these reasons, P-cadherin is considered an important biomarker of poor prognosis in breast cancer (51,52).

In this work, we show that Plk4 overexpression potentiates *anoikis* resistance of non-tumorigenic p53 knock-out mammary epithelial cells, by the induction of a stable hybrid epithelial-mesenchymal phenotype. Moreover, we also show that cells with high Plk4 expression secrete factors that promote *anoikis* resistance of cancer cells in a paracrine way, being P-cadherin one of the mechanistic players in that process. Furthermore, tumors with high expression of Plk4 and P-cadherin are significantly correlated with a worse DFS and OS, as revealed by Kaplan-Meier plots. Therefore, our findings demonstrate that Plk4 overexpression influences the communication between cells and the tumor microenvironment, impacting malignancy in different ways.

## Results

### Plk4 overexpression potentiates *anoikis* resistance and increases colony formation of mammary epithelial cells with a p53 knock-out background

Due to its role in controlling centriole duplication and its deregulation in multiple tumors, participating in tumorigenesis, metastasis and in the chemotherapy response, Plk4 has been put forward as a potential therapeutic target in cancer. However, tumors are known to be very heterogeneous in their constitution. They are composed of distinct pools of cancer cells, including cancer stem cells, non-stem cancer cells and other cells from the tumor microenvironment. Because of the complexity of the tumor microenvironment, the exact molecular mechanism of how Plk4 contributes to tumorigenesis, and its effect in stem-cell features, is still poorly understood.

In the present study, we hypothesized that high levels of Plk4 could contribute to tumorigenesis by inducing stem-like properties, such as *anoikis* resistance, in non-transformed p53-mutated cells, thus potentiating cancer progression.

In order to test this hypothesis, we used the MCF10A-Plk4 cell line, a human mammary non-cancerous cell line, engineered to promote Plk4 overexpression in an inducible manner (53). Upon Doxycycline (Dox) treatment, Plk4 is transiently overexpressed, and consequently CA is promoted. Extra centrosomes affect cells by promoting the formation of multipolar spindles and chromosome misegregation, leading to chromosomal instability and aneuploidy(10,11). Moreover, CA can also be detrimental for cell proliferation, as they can activate the p53 signaling pathway in vertebrate cells, leading to either a G1 cell cycle arrest or apoptosis(54–56). However, cancer cells can usually survive to the presence of multiple centrosomes, because they often do not have a functional p53 pathway(57,58). To ensure cell survival upon centriole number manipulation, and to mimic what occurs in cancer, where the tumor suppressor p53 is often mutated, we successfully knocked out (KO) p53 in the MCF10A-Plk4 cells by using the CRISPR/Cas9 technology (Supplementary fig. 1).

Plk4 overexpression was induced for 24h using 1µg/ml of Dox in MCF10A-Plk4 and MCF10A-Plk4^p53KO^ cell lines. We analyzed *PLK4* mRNA expression by RT-qPCR and, as expected, a significant 4-fold-increase and 9-fold increase were observed after 24h of Dox treatment, in MCF10A-Plk4 and MCF10A-Plk4^p53KO^ respectively, followed by a significant decrease immediately after 24h of Dox removal (Fig. 1A) in both cell lines. Centriole number was quantified in mitotic cells, which normally have a fixed number of 4 centrioles, 2 at each pole of the mitotic spindle (Fig. 1B and Fig. 1C). MCF10A-Plk4^p53KO^ cells showed continuously high percentage of cells with extra centrosomes, even at 96h after Dox removal. However, MCF10A-Plk4 cells presented a sustained decreased of the percentage of cells with centrosome amplification throughout time after Dox removal, equalizing to control levels at 96h (Fig. 1B). The presence of p53 in MCF10A-Plk4 cells might explain why the number of cells with CA decreases after Dox removal, since it is described that, CA activates p53, leading to the elimination of cells with this phenotype(55,59). On the other hand, MCF10A-Plk4^p53KO^ cells with extra centrosomes are maintained for longer in the population (Fig. 1B).

**Figure 1:**
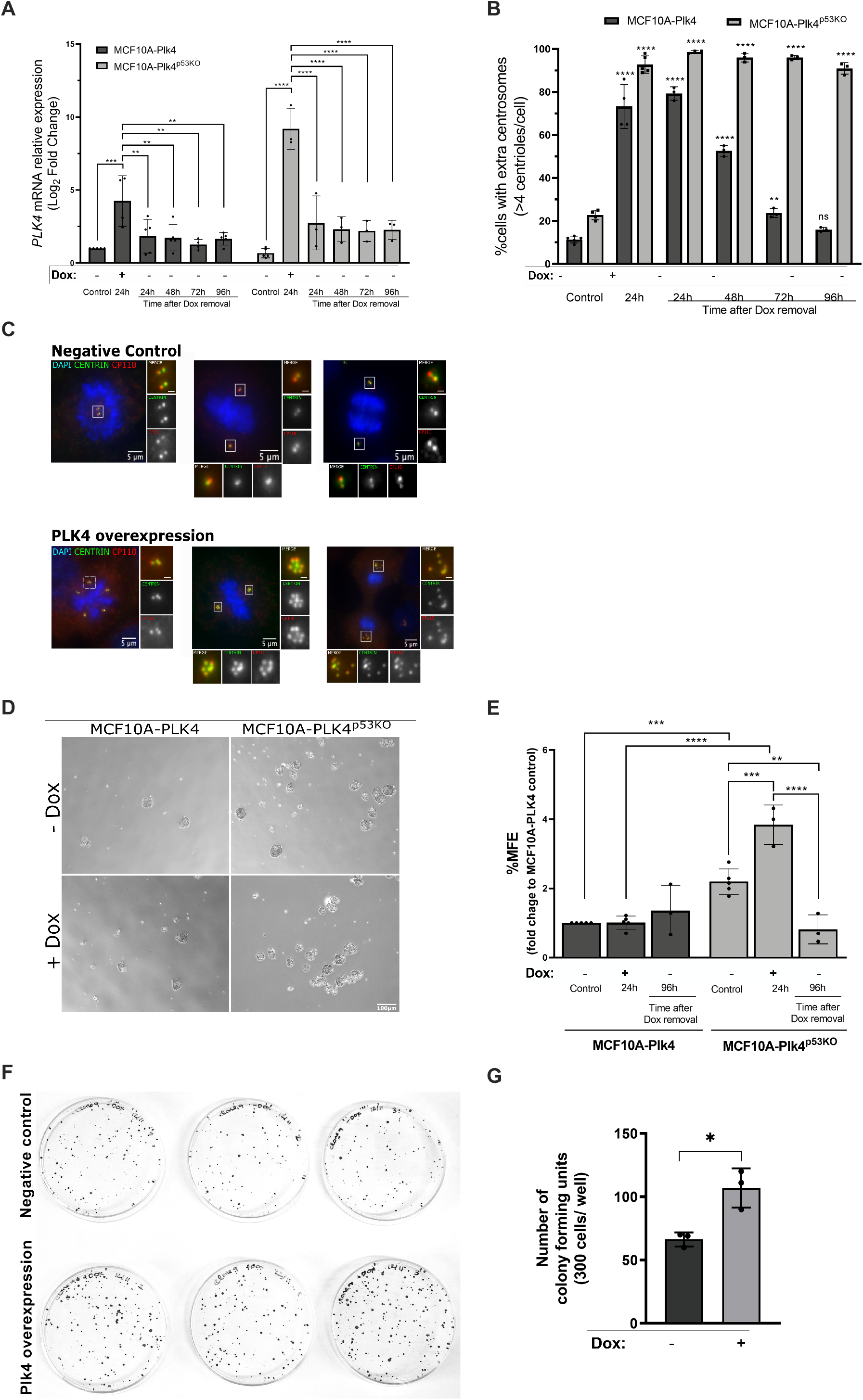
Plk4 overexpression potentiates *anoikis* resistance in MCF10A mammary epithelial cells in a p53 knock-out background. **A)** *PLK*4 mRNA expression levels in MCF10A-Plk4 and MCF10A-Plk4^p53KO^ cells after induction of Plk4 overexpression. After the first stimulus (1µg/ml of Dox for 24h), Dox was removed and cells were kept in Dox-free media, by replenishing media with new fresh media, and collected at different time points (24h, 48h, 72h and 96h) for mRNA analysis. *GAPDH* mRNA expression was used as a housekeeping gene and all conditions were normalized to the control condition (MCF10A-Plk4 control). Data represents average ± SD for at least three independent experiments, p<0.05, One-Way ANOVA test. **B)** Quantification of centriole number in MCF10A-Plk4 and MCF10A-Plk4^p53KO^ mitotic cells 24h of Dox treatment and upon Dox removal. Plk4 overexpression induced centriole amplification in 73% of MCF10A-Plk4 and 92% of MCF10A-PLK4^p53KO^ cells 24h after Dox. 96h following Dox removal, MCF10A-PLK4^p53KO^ maintained high percentage of cells with extra centrosomes, while MCF10A-Plk4 reverted to control levels. To obtain the percentage of cells with >4 centrioles, ≥100 cells were quantified per condition and per experiment. Data represents average ± SD for three independent experiments, and significance to each cell line’s respective control condition. p<0.05, One-Way ANOVA test. **C)** Representative figures displaying cells in negative control conditions (with normal centriole number) and cells with Plk4 overexpression (displaying CA) after Dox treatment. Centrioles were stained with two centriole markers Centrin-1 (green) and CP110 (red). Scale bars: 5µm (main images), 1µm (insets). **D)** Phase contrast images of representative mammospheres formed by MCF10A-Plk4 and MCF10A-PLK4^p53KO^ cells with and without Dox treatment in non-adherent conditions. Magnification: 5X; Scale bar 100µm. **E)** *In vitro* quantification of Mammosphere Forming Efficiency (MFE) of MCF10A-Plk4 and MCF10A-Plk4^p53KO^ cells with and without Plk4 overexpression using 1µg/ml of Dox 24h prior to Mammosphere Forming. MCF10A-Plk4 was used as a control. Moreover, after the first Dox stimulus with 1µg/ml, cells were kept in Dox-free medium for 96h to evaluate MFE. The data is reported as the fold change in the percentage of mammospheres formed/7500 seeded cells ± SD, p<0.05, Ordinary One-Way ANOVA. **F)** Representative image and **G)** quantification of colony forming units of MCF10A-Plk4^p53KO^ cells treated and not treated with 1ug/ml Dox to induce Plk4 overexpression. Plk4 overexpression significantly induces an increase in colony formation in the MCF10A-Plk4^p53KO^ cells. Data shown is average ±SD for three independent experiments, p<0.05, Unpaired t-test

*Anoikis* resistance capacity was assessed by the Mammosphere Forming Efficiency (MFE) assay of MCF10A-Plk4 and MCF10A-Plk4^p53KO^ cell lines. The MFE enables the survival of cells with stem-like features, such as *anoikis* resistance, from single cells, in non-adherent and serum free conditions (60). Only mammospheres equal or greater than 60µm of diameter were counted, as previously described(61). In the control condition, we observed that MCF10A-Plk4^p53KO^ cells formed more mammospheres in comparison to MCF10A-Plk4, suggesting that p53 loss promotes *anoikis* resistance (Fig. 1E). This result was expected since it has been demonstrated that loss of p53 confers *anoikis* resistance capacity to breast cancer cells(31,32,37,62). We also observed that upon Plk4 overexpression in both models, MFE was significantly potentiated, but only in the p53KO background (4-fold increase), suggesting that Plk4 significantly promotes *anoikis* resistance in the absence of p53. Nevertheless, when Dox was removed for 96h, MCF10A-Plk4^p53KO^ cell line lost its capacity to form mammospheres, showing a decreased percentage of MFE. In order to confirm Plk4’s role in inducing carcinogenesis, MCF10A-Plk4^p53KO^ cell line was subjected to the clonogenic assay. Cells were treated/not treated with 1μg/ml Dox to induce Plk4 overexpression and then replated at low cell density (300 cells/plate) and allowed to grow for 10 days. We observed that Plk4 overexpressed cells formed significantly more colonies than non-overexpressed cells (Fig. 1F, G). This result is supported by other studies where Plk4 has been demonstrated to increase colony formation, as well as treatment of Plk4 inhibitor (CFI-400945) significantly reduces cell growth, viability and colony formation in human prostate cancer cells, in multiple embryonal tumor cell lines, and other cancers (41,63–65).

Additionally, we also evaluated whether Plk4 overexpression has an effect on cell viability by taking advantage of the PrestoBlue Cell Viability Reagent. This reagent is reduced by metabolically active cells, providing a quantitative measurement of viable and proliferating cells. Interestingly, the knock-out of p53 did not alter cell viability of MCF10A-Plk4^p53KO^ when compared to the parental cell line MCF10A-Plk4 (Supplementary fig. 2). By treating cells with 1μg/ml of Dox for 24h to induce Plk4 overexpression, we observed that it significantly altered cell viability when compared to the control condition in both cell lines. However, cell viability of MCF10A-Plk4^p53KO^ was significantly higher than in MCF10A-Plk4, when Plk4 was overexpression was induced by Dox. These results demonstrate that, besides contributing to *anoikis* resistance, Plk4 overexpression (in the p53KO background) also potentiates cell viability, demonstrating its putative role in tumorigenesis.

As Plk4’s effect in *anoikis* resistance might be specific to the MCF10A epithelial cell line, we therefore explored Plk4’s role in *anoikis* resistance in the RPE cell line. The RPE cell line is an hTERT-immortalized retinal pigment epithelial cell line, which we engineered to enable the inducible expression of Plk4 upon doxycycline (Dox) treatment (RPE-Plk4 cell line). To confirm that Plk4 is overexpressed upon Dox treatment, cells were treated for 24h with 1μg/ml of Dox, and *PLK4 mRNA* levels and centrosome number was quantified. We observed that, upon Plk4 overexpression, the RPE-Plk4 cell line presented a significant 4-fold increase in *PLK4* mRNA levels (in comparison to not treated condition) (Supplementary fig. 3A). Moreover, we also observed an increase from 5% (not treated condition) to 92% of mitotic cells with >4 centrioles upon Plk4 overexpression (Supplementary fig. 3B), confirming that this cell line is responsive to Dox treatment inducing Plk4 overexpression. After, we silenced p53 expression in the RPE-Plk4 cell line, and performed spheres assay upon Dox-treatment to induce Plk4 overexpression. As it can appreciate in Supplementary fig. 3C, RPE-Plk4 cells were able to form spheres, but only in the condition where p53 expression was silenced. Moreover, when Plk4 was overexpressed upon Dox treatment, the sphere efficiency of RPE-Plk4 was potentiated, demonstrating that Plk4 overexpression, in the absence of p53, induces *anoikis*-resistance capacity of normal epithelial cells. These findings support our previous results in the MCF10A cell line, showing that Plk4 contributes to tumorigenesis by increasing the ability of p53 knock-out mammary epithelial cells to circumvent *anoikis*.

### The impact of Plk4 overexpression on *anoikis* resistance is independent of its role in centrosome amplification

Since we observed that, at 96h following Dox removal, MCF10A-Plk4^p53KO^ cells maintained a high percentage of CA, but low levels of Plk4, we wondered whether *anoikis* resistance capacity of these cells was being mediated by CA or by Plk4 overexpression. To tackle this question, we used a cell line with a truncated form of Plk4 (Plk4^1-608^), which retains kinase activity, but does not induce CA, when Plk4 is transiently overexpressed(53,66). To confirm the increase in Plk4 levels, but not in centrosome number after Dox treatment in MCF10A-Plk4^1-608^ cells, we analyzed *PLK4* mRNA levels (Fig. 2A) and centrosome number (Fig. 2B) after overexpressing Plk4 by using 1µg/ml of Dox. As appreciated in Figure 2A, a significant 3.5-fold increase in Plk4 mRNA levels was observed in MCF10A-Plk4^1-608^ cells upon 24h of Dox, while, as expected, no CA was observed (Fig. 2B). We next asked whether the observed *anoikis* resistance was due to increased Plk4 levels or to CA. In order to test this, MCF10A-Plk4^1-608^ cells was subjected to the MFE assay. Interestingly, when Plk4 is overexpressed, MCF10A-Plk4^1-608^ was able to form more mammospheres, when compared to control conditions, suggesting that Plk4 catalytic activity, promotes *anoikis* resistance, independently of its role in CA (Fig.2C).

**Figure 2:**
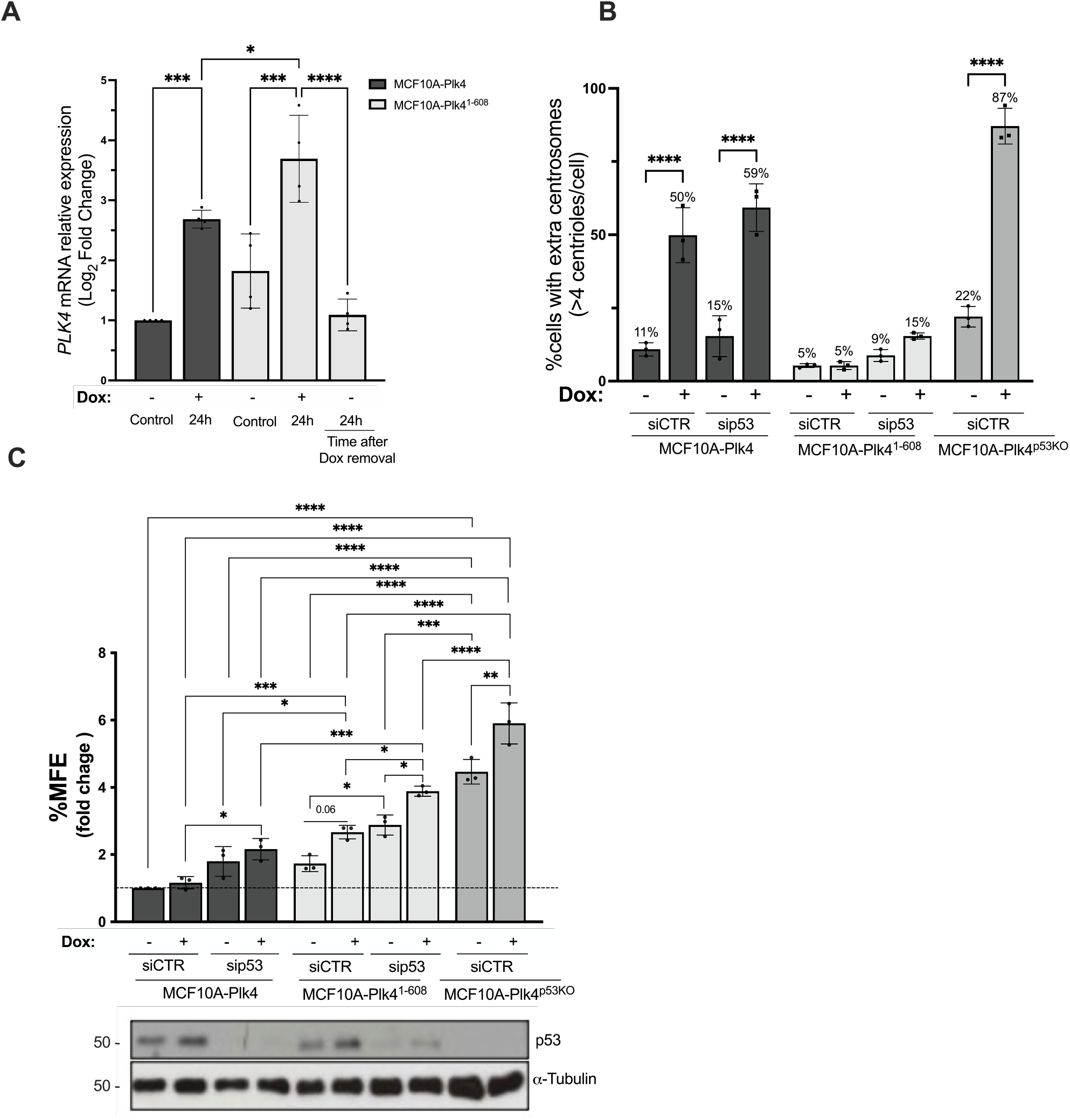
Plk4 catalytic activity is involved in *anoikis* resistance of mammary epithelial cells, independently of its role in centrosome amplification. **A)** *PLK*4 mRNA expression levels in MCF10A-Plk4^1-608^ cells after induction of Plk4 overexpression using 1µg/ml of Dox for 24h. After this stimulus, Dox was removed by replenishing media with new fresh media and collected 24h after Dox removal for mRNA analysis. *GAPDH* mRNA expression was used as a housekeeping gene and all conditions were normalized to the MCF10A-Plk4 control condition. Data represents average ± SD for at least three independent experiments, p<0.05, One-Way ANOVA test. **B)** Quantification of centriole number in MCF10A-Plk4, MCF10A-Plk4^p53KO^ and MCF10A-Plk4^1-608^ mitotic cells 24h after Plk4 overexpression in silenced p53 (sip53) and not silenced (siCTR) conditions. Plk4 overexpression induced centriole amplification in 50% of MCF10A-Plk4 and 87% of MCF10A-PLK4^p53KO^ cells, but not in MCF10A-Plk4^1-608^ (5%). When Plk4 is overexpressed in a p53 silenced background, MCF10A-Plk4 showed an increase of 59% of cells with extra centrosome, and MCF10A-Plk4^1-608^ presented a 15% increase of CA, in comparison to the respective controls. To obtain the percentage of cells with >4 centrioles, ≥100 cells were quantified per condition and per experiment. Data represents average ± SD for three independent experiments, and significance to each cell line’s respective control condition. p<0.05, One-Way ANOVA test. **C)** *In vitro* quantification of Mammosphere Forming Efficiency (MFE) assay of MCF10A-Plk4, MCF10A-Plk4^p53KO^ and MCF10A-Plk4^1-608^ cells with and without Plk4 overexpression using 1µg/ml of Dox 24h prior to the assay with and without p53 silencing. MCF10A-Plk4 was used as a control. The data is reported as the fold change in the percentage of mammospheres formed/7500 seeded cells ± SD, p<0.05, Two-Way ANOVA.

As previously mentioned, loss of p53 confers *anoikis* resistance capacity to breast cancer cells. Thus, we finally asked whether after p53 removal, MCF10A-Plk4 and MCF10A-Plk4^1-608^ cells would have a higher ability to form mammospheres. By silencing p53 with a specific siRNA (Fig. 2C), both cell lines showed an increase in their ability to resist to *anoikis*, and this effect was significantly potentiated when Plk4 was overexpressed upon Dox treatment (Fig. 2C). Furthermore, we observed that in the absence of p53, MCF10A-Plk4 and MCF10A-Plk4^1-608^ cell lines presented a small increase in the percentage of cells with extra centrosomes after Dox treatment (Fig. 2B). These results reinforce the role of p53 in Plk4-induced *anoikis* resistance, as well in the control of centrosome number.

### Plk4 overexpression induces the acquisition of a hybrid E/M phenotype on MCF10A-Plk4 p53KO mammary epithelial cells

Loss of p53 by epithelial cells has been described to confer stem-like properties and to induce EMT, allowing them to circumvent *anoikis*, contributing to tumor progression(31–33,67). Thus, we went to explore the EMT profile of MCF10A-Plk4 and MCF10A-Plk4^p53KO^ cells. Non-induced MCF10A-Plk4 cells showed a clear epithelial phenotype, with high levels of E-cadherin and low expression of mesenchymal markers, such as Vimentin, Snail/Slug and Zeb2 (Fig. 3). On the other hand, MCF10A-Plk4^p53KO^ cell line exhibited a mesenchymal phenotype, with low E-cadherin expression, and high levels of Vimentin and Snail/Slug. However, when Plk4 was induced in the p53KO context, there was a reversion of the mesenchymal phenotype to a hybrid EMT phenotype, with an increase in the expression of E-cadherin, as well as of the EMT transcription factors Snail/Slug and Zeb2, and a concomitant decrease in Vimentin levels (Fig. 3). Altogether, these results indicate that Plk4 overexpression, in a p53KO context, induces the stabilization of a hybrid EMT phenotype in non-tumorigenic mammary epithelial cells.

**Figure 3:**
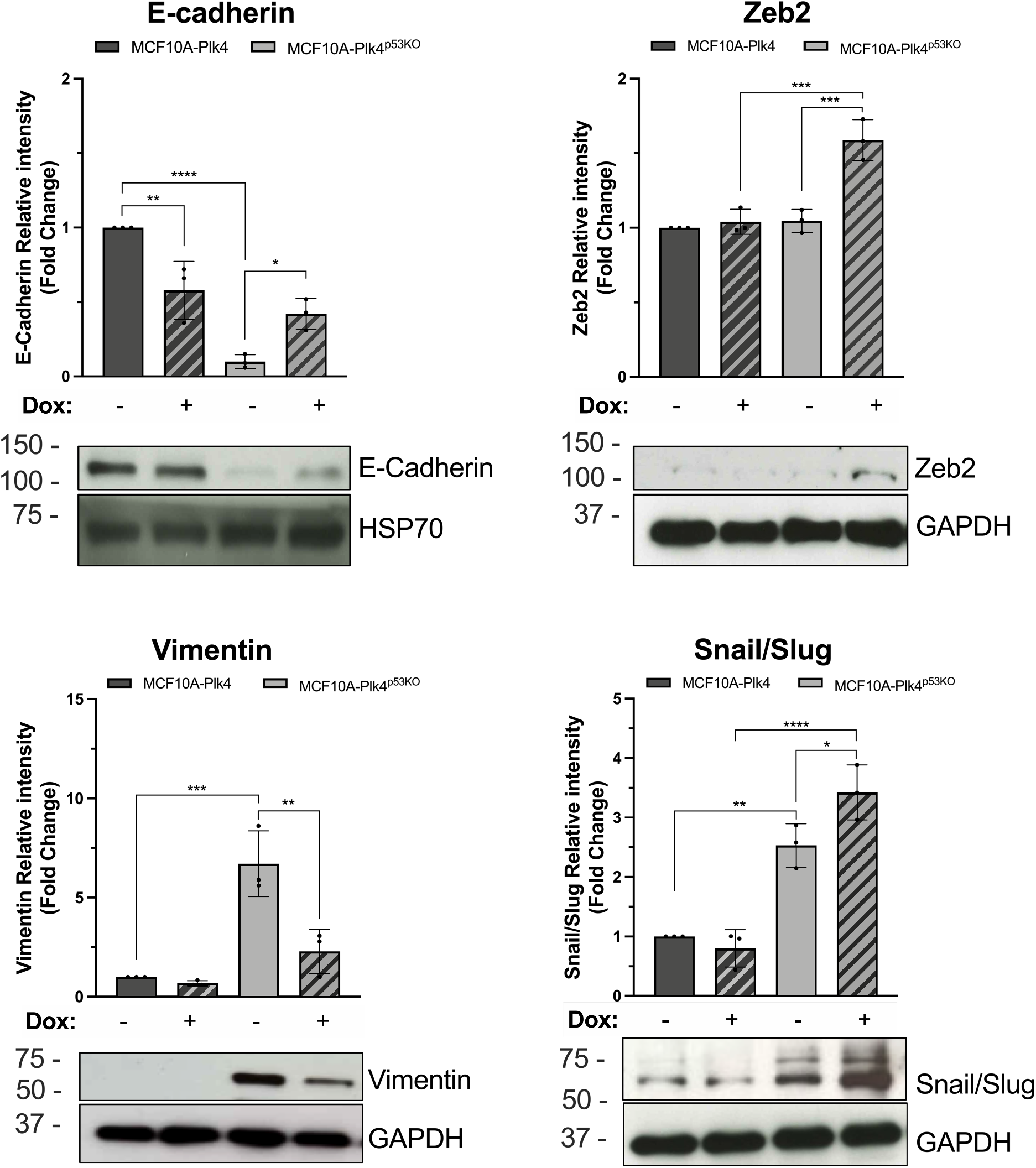
MCF10A-Plk4 p53KO mammary epithelial cells with centrosome amplification show a hybrid epithelial-mesenchymal phenotype. Western Blot analysis of MCF10A-Plk4 and MCF10A-Plk4^p53KO^ cells with and without Plk4 overexpression using 1μg/ml of Dox for 24h. Representative western blot images and quantification of their relative integrated intensity of Epithelial (E-cadherin) and Mesenchymal markers (Vimentin, Zeb2, Snail/Slug). MCF10A-Plk4 cells presents an epithelial profile, with high levels of E-cadherin and low levels of Zeb2, Vimentin and Snail/Slug. On the other hand, the p53KO cells showed a mesenchymal profile, with low expression of E-cadherin and increased expression of Vimentin and Snail/Slug. When Plk4 is overexpressed, p53KO cells acquire a Hybrid EMT phenotype, with high Zeb2, Snail/Slug and E-cadherin expression. Hsp70 (for E-cadherin) and GAPDH were used as loading controls. Integrated intensity was measured by ImageJ Software (National Institutes of Health). All conditions were normalized to the control condition (MCF10A-Plk4, non-induced). Data shown is the fold change in pixels integrated intensity ± SD for three independent experiments, p<0.05, Ordinary One-Way ANOVA.

In neuroblastoma, Plk4 promotes EMT through PI3K/Akt signaling pathway, by downregulating E-cadherin and upregulating EMT-related factors, such as Snail/Slug(25). On the other hand, Plk4 also promotes EMT via Wnt/*β*-catenin in colorectal cancer(41). Interestingly, in esophageal squamous cell carcinoma, Plk1 overexpression, another member of the polo-like kinase family, has been demonstrated to trigger *anoikis* resistance through regulation of *β*-catenin expression, by directly binding to the NF-kB subunit RelA, which inhibits the ubiquitination and degradation of *β*-catenin(68). Importantly, NF-kB signaling pathway, which is activated upon cellular stress, can regulate the expression of genes required for centrosome duplication, being Plk4 a direct NF-kB target gene(69). Studies, have also documented a crosstalk between NF-kB and p53(70). It is important to point out that p53 has been shown to downregulate Plk4 and Plk1 expression(71,72). These data suggests that regulation of these kinases provides another rout of potential crosstalk within cells potentiating cancer progression. However, the exact mechanism through which Plk4 might be mediating a hybrid EMT, protecting cells against *anoikis* is still poorly explored and needs further study.

### P-cadherin expression is increased in Plk4 induced-MCF10A-PLK4^p53KO^ cells and is required for *anoikis* resistance in an autocrine and paracrine way

Recently, the concept of hybrid EMT has emerged and has been shown to be extremely important for cancer cells to become invasive and metastatic in a collective manner(73,74). We have previously demonstrated that overexpression of the cell-cell adhesion molecule P-cadherin promotes collective cell invasion, stem cell properties and tumorigenesis(43,44,50,61). Moreover, P-cadherin expression is early promoted by EMT-inducers, such as hypoxia, driving *anoikis* resistance in breast cancer cells(44) and being a putative biomarker and stability factor of a hybrid EMT phenotype(43,52).

Therefore, we analyzed P-cadherin mRNA and protein expression in both MCF10A-Plk4 and MCF10A-Plk4^p53KO^ cells. Interestingly, we observed that P-cadherin is mainly expressed by MCF10A-Plk4^p53KO^ cells (Fig. 4A, 4B), and its expression is significantly increased, both at mRNA and protein levels, when Plk4 is overexpressed. Nevertheless, no alterations in P-cadherin expression were observed in MCF10A-Plk4 cells, where p53 is normally expressed.

**Figure 4:**
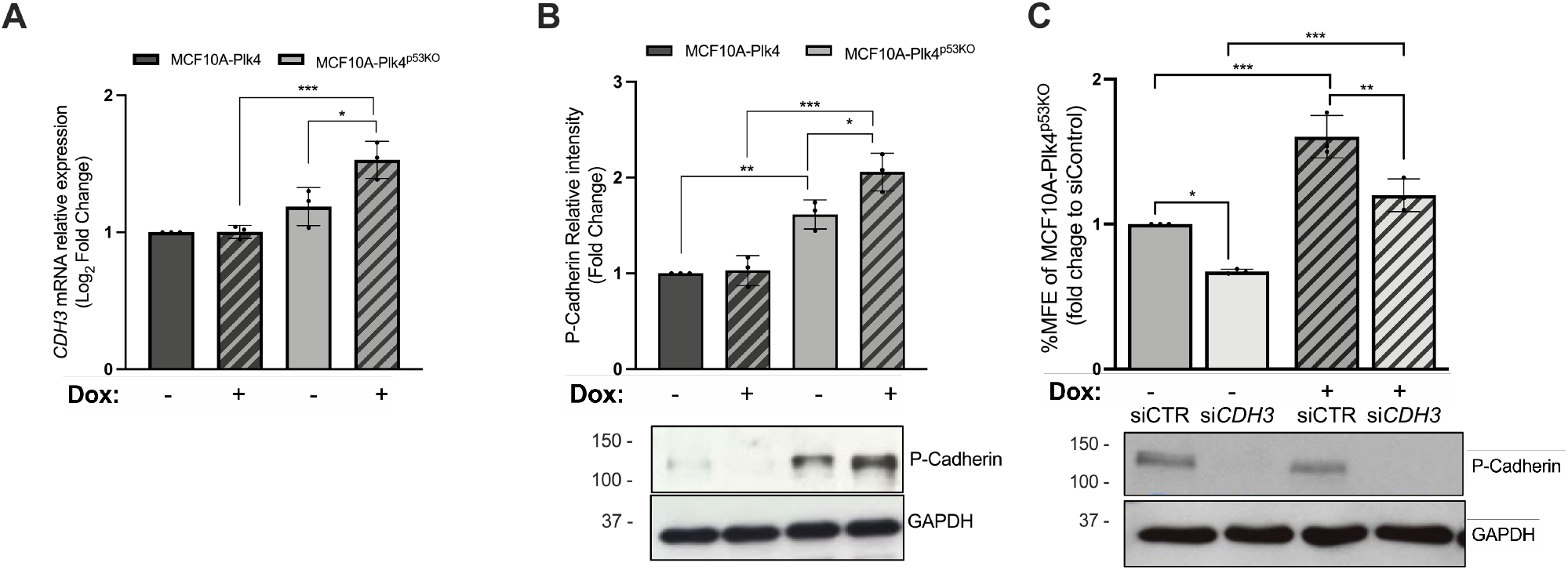
P-cadherin expression is increased in Plk4 induced-MCF10A-Plk4^p53KO^ cells and is partially required for resistance to *anoikis*. P-Cadherin expression is increased in p53KO cells treated with doxycycline, but not observed in MCF10A-Plk4, both at the **A)** (*CDH3*) mRNA and **B)** at the protein level. P-cadherin gene (*CDH3*) expression was assessed by qRT-PCR and protein levels by Western Blot of MCF10A-Plk4 and MCF10A-Plk4^p53KO^ cell lines treated and not treated with doxycycline. All conditions were normalized to the MCF10A-Plk4 control condition. Data shown is average ± SD for three independent experiments, p<0.05, Ordinary One-Way ANOVA. **C)** Silencing of P-cadherin in Plk4-induced MCF10A-Plk4^p53KO^ significantly decreased their capacity to form mammospheres. *CDH3* gene silencing was performed in MCF10A-Plk4^p53KO^ cells using a validated siRNA, specific for *CDH3*. A scrambled siRNA with no homology to any gene, was used as a negative control (siCTR). Following siRNA transfection, cells were treated or not treated with 1μg/ml of Dox for 24h. MFE was performed after Dox treatment. All conditions were normalized to siControl. Western Blot bellow represents the validation of P-Cadherin silencing. Data shown is average ± SD for three independent experiments, p<0.05, Ordinary One-Way ANOVA.

Based on these observations, we next evaluated whether P-cadherin expression was required for the Plk4-induced *anoikis* resistance in p53KO cells. Thus, *CDH3*/P-cadherin was silenced in MCF10A-Plk4^p53KO^ cells using specific siRNA for the *CDH3* gene. A scrambled siRNA, with no homology to any gene, was used as a negative control (siCTR). Following siRNA transfection, cells were treated or not treated with 1μg/ml of Dox for 24h (siCTR+Dox and si*CDH3*+Dox) and MFE assay was performed. As expected, and previously shown, by inducing Plk4 overexpression with Dox, cells formed significantly more mammospheres in comparison to non-treated cells (Fig. 4C). Interestingly, the silencing of P-cadherin expression in the MCF10A-Plk4^p53KO^ cells significantly decreased their capacity to form mammospheres, independently of Plk4 overexpression. However, the percentage of MFE observed upon si*CDH3* and induction of Plk4 in MCF10A-Plk4^p53KO^ cells did not reach the MFE percentage observed without induction, whether *CDH3* was silenced or not. This suggests that Plk4 overexpression is able to induce *anoikis* resistance in p53KO MCF10A cells, and this effect is partially mediated by P-cadherin expression.

A non-cell autonomous role for CA has been previously reported(75,76), where cells with extra centrosomes caused by Plk4 overexpression induce paracrine invasion of other cells by secreting extracellular vesicles with pro-invasive factors. Since there is differential protein secretion in cells with centrosome amplification via Plk4 overexpression, we decided to investigate whether high Plk4 expression would also play a role in potentiating paracrine *anoikis* resistance of cancer cells in the tumor microenvironment. Thus, BT20 breast cancer cells were treated with the conditioned media of MCF10A-Plk4^p53KO^ cells for 48h and performed the MFE assay. We observed a significant increase on the capacity of these cancer cells to form mammospheres when treated with the conditioned media of Plk4-induced MCF10A^p53KO^ (Fig. 5A) when compared to control condition. Because the increased mammosphere capacity in BT20 might be due to BT20’s exposure of a CM derived from a different cell line, rather than due to Plk4 overexpression, we also treated MCF10A-Plk4^p53KO^ cells (CM receiver) with the CM of Plk4 overexpressed MCF10A-Plk4^p53KO^ cells (CM donor) for 48h and measured their MFE (supplementary fig. 4). We observed that MCF10A-Plk4^p53KO “^CM receiver” cells formed significantly more mammospheres than control (not treated condition) and serum-free condition (CM was produced in serum-free media). Interestingly, the CM from MCF10A-Plk4^p53KO^ “donor” cells without Plk4 overexpression did not induce an increase in mammosphere formation in MCF10A-Plk4^p53KO “^receiver” cells, suggesting that this effect is mainly mediated by Plk4 overexpression.

**Figure 5.**
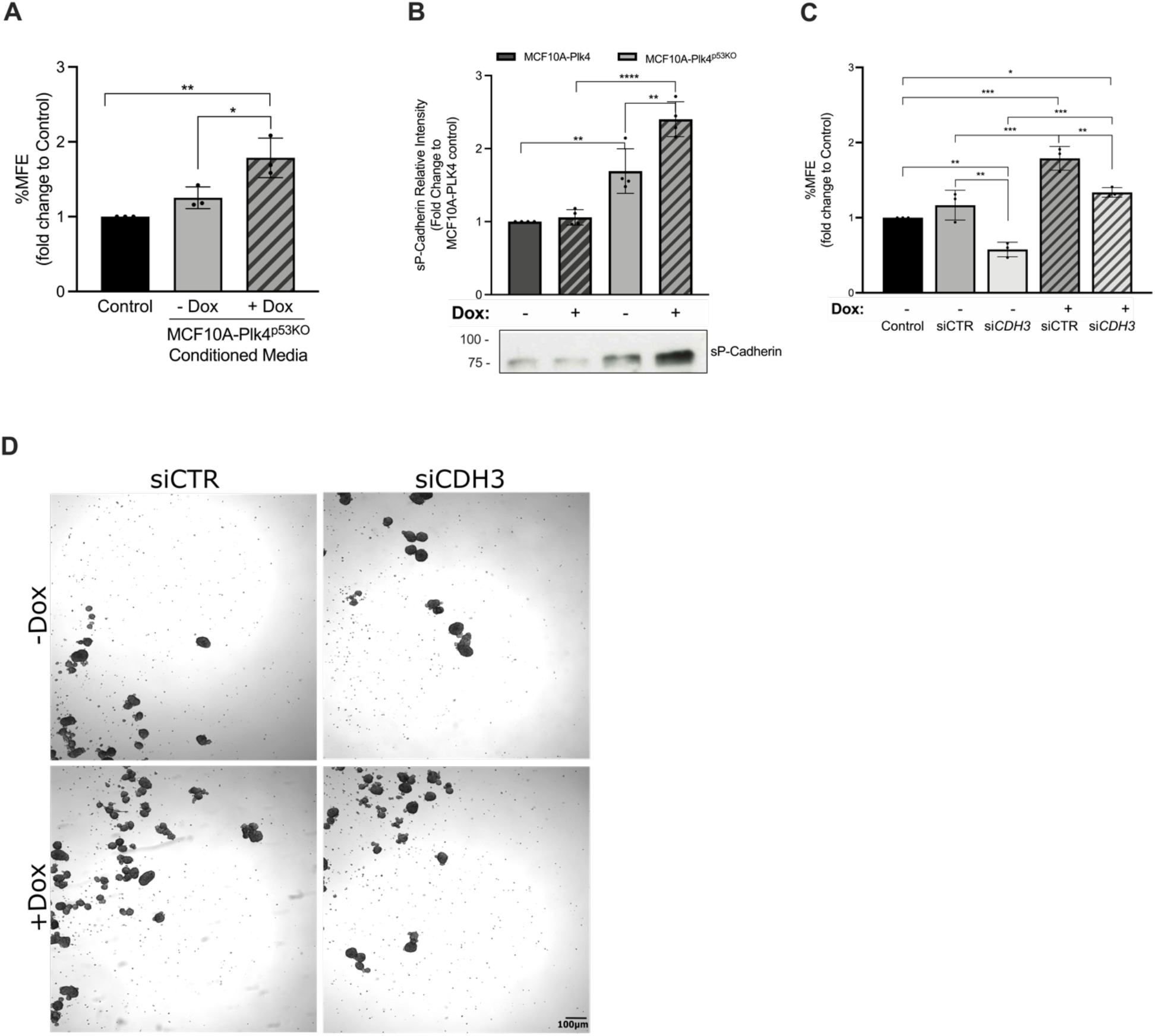
Soluble P-Cadherin (sP-cad) is partially required for *anoikis* resistance of breast cancer cells mediated through the conditioned media of p53KO cells with centrosome amplification. **A)** MFE assay of BT20 breast cancer cells under the following conditions: (control) nontreated; (Dox-) treated with conditioned media from MCF10A-Plk4^p53KO^ without Plk4 OE; (Dox+) treated with conditioned media from MCF10A-Plk4^p53KO^ with Plk4 OE. There is a significantly increase MFE when BT20 cells were treated with the conditioned media of MCF10A-Plk4^p53KO^ with induced Plk4 overexpression. All conditions were normalized to the control condition (BT20 control). Data shown is average ± SD for three independent experiments, p<0.05, Ordinary One-Way ANOVA. **B)** Western blot analysis of conditioned media from MCF10A-Plk4 and p53KO cell line showing different expression levels of sP-cad. P-cadherin ectodomain cleaved (sP-cad) expression is increased in p53KO cells treated with Dox. MCF10A-Plk4 cells have lower P-cadherin basal levels, which is not affected by Plk4 overexpression. Data shown is average ± SD for four independent experiments, p<0.05, Ordinary One-Way ANOVA. **C)** MFE assay of BT20 breast cancer cells incubated with the following conditioned media: (Control) no treatment; (siCTR; Dox-) MCF10A-PLK4^p53KO^-siCTR without Plk4 OE; (siCDH3; Dox+) MCF10A-Plk4^p53KO^ depleted of P-cadherin without Plk4 OE; (siCTR; Dox+) MCF10A-Plk4^p53KO^-siCTR with Plk4 OE; (siCDH3; Dox+) MCF10A-Plk4^p53KO^ depleted of P-cadherin with Plk4 OE. P-cadherin-silencing significantly affects mammosphere forming capacity. Data shown is average ± SD for three independent experiments, normalized to BT20 control, p<0.05, Ordinary One-Way ANOVA. **D)** Phase contrast images of representative mammospheres formed by BT20 breast cancer cells treated with MCF10A-Plk4^p53KO^’s conditioned media of siCTR and siCDH3 cells with and without Dox treatment in non-adherent conditions. Magnification: 2X; Scale bar 100µm.

We have previously shown that P-cadherin overexpression in breast cancer cells promotes an increase in cell migration and invasion due to the secretion of pro-invasive factors, such as MMP1 and MMP2, which then leads to P-cadherin ectodomain cleavage (soluble Pcad, sP-cad) that also has pro-invasive activity by itself(49). Given that the conditioned media of MCF10A-Plk4^p53KO^ cells increased *anoikis* resistance of cancer cells, we investigated if the soluble/cleaved P-cadherin form (sP-cad) was also being secreted by these cells. We found that, sP-cad expression was significantly increased in MCF10A-Plk4^p53KO^ cells (not induced condition), which was not observed in MCF10A-Plk4 (Fig. 5B). Importantly, sP-cad expression was significantly potentiated when Plk4 was induced in the MCF10A-Plk4^p53KO^, suggesting that Plk4 plays a direct role in sP-cad expression in the p53KO context. Finally, we went to evaluate if sP-cad was also required for the induction of paracrine *anoikis* resistance of breast cancer cells, by treating BT20 breast cancer cells with the conditioned media of MCF10A-Plk4^p53KO^ upon *CDH3* silencing. We observed a significant 0.5-fold decrease in the ability of BT20 cells to form mammospheres when treated with conditioned media of MCF10A-Plk4^p53KO^, with *CDH3* silencing, independently of Plk4 overexpression (Fig. 5C). Moreover, we also observed that cancer cells treated with the conditioned media of cells with both Plk4-overexpressed and silenced P-cadherin presented a decreased capacity to form mammospheres, but they still have higher MFE than control cells. This emphasizes our previous result, where high levels of Plk4 induces *anoikis* resistance partially depending on P-cadherin, also in a paracrine way.

Overall, our results highlight the role of Plk4 and P-cadherin, in influencing the communication between different cells to promote malignancy.

## Discussion

Dysregulation of Plk4 has been found in several types of cancers and shown to cause loss of centrosome numerical integrity, promoting genomic instability. Furthermore, increased Plk4 expression has been linked to cancer metastasis and to chemotherapy resistance. However, the control of centriole duplication may not be the only relevant function of Plk4 in carcinogenesis.

Herein, we investigated if Plk4 could play a role in carcinogenesis, specifically by inducing stem-like features, such as *anoikis* resistance. We demonstrated that loss of the tumor suppressor p53 is necessary to maintain Plk4 induced-CA throughout time. Moreover, we also showed that Plk4 overexpression promotes *anoikis* resistance in non-tumorigenic mammary epithelial cells, in a p53 knockout background. As centrosome amplification is a direct cause of Plk4 overexpression, CA could also be contributing to *anoikis* resistance. By taking advantage of a cell line with truncated form of Plk4 (Plk4^1-608^) that retains kinase activity but does not induce centrosome amplification when Plk4 is overexpressed, we observed that these cells are able to resist *anoikis* when Plk4 is overexpressed. These results demonstrates that the *anoikis* resistance observed is directly mediated by Plk4 kinase activity, independently of its role in promoting CA. Moreover, we also showed that p53-silenced Plk4^1-608^ cells are more resistant to *anoikis* than p53 WT Plk4^1-608^ cells, which is then potentiated when Plk4 is overexpressed. The same was observed by transiently inducing Plk4 in the MCF10A-Plk4^p53KO^, where cells became *anoikis* resistant, confirming that Plk4 potentiates *anoikis* resistance in the p53 knock-out background. Moreover, we also demonstrated that in the MCF10A-Plk4^p53KO^ cell line, Plk4 overexpression significantly increases cell viability and colony formation, reinforcing its putative role in tumorigenesis.

Loss of p53 has been described to confer stem-like properties, to potentiate *anoikis* resistance of cells, and to induce an epithelial to mesenchymal transition (EMT)(31,32). In this work, we showed that high Plk4 levels in a p53KO context leads to a hybrid EMT phenotype, with increased expression of P-cadherin. In fact, *anoikis* resistance and hybrid EMT phenotype are crucial features for cancer progression and metastatic colonization(36,77,78). In accordance to other studies, in which P-cadherin has been shown to be promoted by EMT-inducers, such as hypoxia, driving *anoikis*-resistance capacity in breast cancer cells(43,44), we demonstrate that P-cadherin expression is partially required for *anoikis* resistance observed in the MCF10A-Plk4^p53KO^ cells, as well as in breast cancer cells exposed to their conditioned media.

Previous reports have shown that the hybrid EMT state is critical to stemness, independently of phenotypic plasticity(42). Both epithelial and mesenchymal traits need to be co-expressed within an individual cancer cell for efficient tumorigenicity, as mesenchymal properties are important for the intravasation from the primer tumor and survival in blood circulation, whereas epithelial traits are essential for the metastatic colonization of distant organs(42,79). P-cadherin has been demonstrated as a promising biomarker of the hybrid EMT state, based on the rationale that P-cad expression disturbs epithelial cell-cell adhesion and promotes the acquisition of a more undifferentiated cell phenotype, giving these cells a phenotypic state between epithelial and mesenchymal morphology(43,61). Moreover, cells overexpressing P-cadherin show increase therapy resistance, stem cell properties, a more aggressive and invasive behavior(43). These properties point to a hybrid EMT state, with increased plasticity and metastatic capacity. In fact, P-cadherin has been described as a poor prognostic biomarker in basal-like breast tumors, and to correlate with histological grade, being essentially present in high-grade tumors(45,51,52). Interestingly, overexpression of Plk4 has also been shown to have a prognostic value associated with worse DFS and OS of breast cancer patients(80,81). Moreover, unpublished data from our lab shows that P-cadherin expression is significantly associated to the expression of the EMT transcription factor Zeb2. Zeb2 has also been pointed out as possible hybrid-EMT marker in breast cancer, as its mRNA is highly expressed in cells with hybrid E/M and Mesenchymal phenotypes (82). In accordance, we also showed that in the absence of p53, Plk4 overexpression induces an increase in Zeb2 expression, reinforcing the role of Plk4 in inducing a hybrid EMT phenotype.

By taking advantage of the publicly available data from Kaplan Meier plotter (https://kmplot.com/analysis/), where RNA seq data is available for n=2976 breast cancer patients, we observed that patients with tumors co-expressing high Plk4 and P-cadherin expression show a worse prognosis (Supplementary fig. 5). Besides breast cancer, Plk4 has been found to be upregulated in most solid tumors, such as gastric, pancreatic, lung, melanoma, cervical, osteosarcoma and brain (neuroblastoma, glioblastoma, medulloblastoma) tumors, and associated with shortened patient survival in these tumors(18,80,81).

In conclusion, our observations have important implications for understanding the potential link between Plk4 levels, cancer and tumor microenvironment, as well as clinical outcomes. Our data shows that high Plk4 expression is associated with malignant features such as *anoikis* resistance and EMT in cancerous and non-cancerous cells in the tumor microenvironment, endorsing its role in tumor initiation and/or progression. As emerging data has been supporting the idea that Plk4 plays an important role in tumorigenesis, more comprehensive research on the exact signaling pathways and on the present and next-generation Plk4 inhibitors, following successful clinical studies, may provide a new dimension for innovative cancer therapies.

## Materials and methods

### Cell culture and growth conditions

Cell lines were maintained at 37°C with humidified 5% CO_2_ atmosphere.

Human mammary epithelial MCF10A-Plk4 and MCF10A-Plk4^1-608^ cells (kind gift from Susana Godinho, from Barts Cancer Institute, Queen Mary University of London) and MCF10A-Plk4-p53 knock-out cells were grown in DMEM/F12 media supplemented with 5% Horse Serum, 20ng/ml epidermal growth factor (EGF), 10μg/ml insulin, 100ng/ml cholera toxin, 0.5μg/ml hydrocortisone, 100U/ml penicillin and streptomycin.

BT20 cancer cells were grown in DMEM media supplemented with 10% fetal bovine serum (FBS), and 100U/ml penicillin and streptomycin.

RPE-Plk4 cells were grown in DMEM/F12 media supplemented with 10% Tet-free fetal bovine serum (FBS), and 100U/ml penicillin and streptomycin.

### P53 Knock-out cell line

For the generation of p53 knock-out (p53KO) cell line, CRISPR/Cas9 plasmid (sc-416469) and p53 Homologous direct Recombinant-HDR plasmids (sc-416469-HDR) were transfected into MCF10A-Plk4 cells using the Neon® Transfection system. After transfection recovery, cells were selected with puromycin in order to select p53KO stable cell line. To have monoclonal stable p53KO clones, puromycin resistant cells were sorted into single cell clones by using a BD FACS Aria IIu.

To validate if p53 was successfully knocked-out, cells were exposed to three different conditions to determine the functionality of the p53 protein: treatment with 1 and 3µM of Doxorubicin (mild and severe DNA damage, respectively) for 4h and kept for 24h in drug free medium and, a third condition where 1µg/ml of Doxycycline was added to cells for 24h (inducing extra centrosomes). After the 24h, cells were harvested for western-blot analysis. Total protein levels of p53 and p21, a downstream target of p53, was assessed by Western Blot.

### Plk4 overexpression - Doxycycline (Dox) treatment

Cells were grown until 80-85% confluence and for every experiment, cells were treated with 1µg/ml of Dox (Merck, Darmstadt, Germany) to induce Plk4 overexpression for 24h. After 24h of treatment, cells were washed twice with 1X phosphate buffer solution (PBS), trypsinized and counted, before starting any specific assay. In parallel, a control condition (no doxycycline) was also performed.

### Immunofluorescence

Adherent cell lines were grown on glass coverslips (VWR, #631-0150) and fixed using cold methanol for 10 min at -20 °C. After fixation, cells were washed with 1X PBS and incubated for 30min at room temperature (RT) with 1X PBS containing 10% FBS. Afterwards, cells were incubated for 1h30min at RT with primary antibodies Centrin-1(20H5) (1/1000, Merck 04-1424) and CP110 (1/250, Jiang et al, 2012(83)) diluted in 1X PBS + 10%FBS. After the incubation with the primary antibodies, cells were washed with 1X PBS three times, 10 min each. Cells were then incubated for 1h at RT with corresponding secondary antibodies Alexa Fluor 488 (Alfagene, #A11034) and Alexa Fluor 549 (Alfagene, #A11032), diluted at 1:500 in 1X PBS with 10%FBS. After the secondary antibodies’ incubation, cells were washed four times with 1X PBS. DAPI (1:500) was added on the second wash to stain DNA. The coverslips were then mounted on slides using DAKO Faramount Aqueous Mounting Medium (Agilent, #S302580-2). The slides were kept for 24h to allow the mounting media to cure before use.

Two centriolar markers were always used, in order to avoid false positive. Only structures positive for the two markers were considered as centrioles.

### Image acquisition and centrosome quantification

Mitotic cells were observed using a Leica DMI6000 (Leica Microsystems, Germany) Microscope and images were acquired with a Hamamatsu FLASH4.0 (Hamamatsu, Japan) camera, using the HCX PL APO CS 63x/1.30 GLY 21°C objective, and the LAS X Software. Images were taken in Z-Stacks in a range of 10-14*μ*m, with a distance between planes of 0.2*μ*m.

We consider as centriole amplification, when mitotic cells presented more than four (>4) centrioles. In order to obtain the percentage of cells with extra centrioles, at least 100 cells were analyzed for centriole number per condition and per experiment and only centrioles positive for the two centriolar markers (Centrin-1 and CP110) were considered. Centrosomes were quantified manually, using the Fiji/Image J Software (National Institutes of Health).

### Mammosphere/Sphere forming efficiency (MFE) assay

Cells were enzymatically harvested and manually disaggregated to form a single-cell suspension and resuspended in cold PBS. Cells were then plated at the density of 750cells/cm^2^ in non-adherent culture conditions, in 6-well plates coated with 1.2% poly (2-hydro-xyethylmethacrylate)/95% ethanol (Sigma-Aldrich) and allowed to grow for 5 days, in DMEM/F12 containing B27 supplement (Invitrogen), 500 ng/mL of hydrocortisone, 40 ng/mL insulin, 20 ng/ mL EGF in a humidified incubator at 37°C and 5% (v/v) CO_2_. MFE was calculated as the number of mammospheres (≥60 μm) formed divided by the number of cells plated, and presented as percentage.

### Protein extraction and western blot analysis

Protein extracts from cultured cells were prepared by using catenin lysis buffer [1%(v/v) Triton X-100 and 1%(v/v) NP-40 (Sigma-Aldrich, USA) in deionized phosphate-buffered saline (PBS)] supplemented with 1:7 proteases inhibitors cocktail (Roche Diagnostics GmBH, Germany) for 10 min, at 4 °C. Cell lysates were vortexed and centrifuged at 14000rpm at 4°C, for 10min. Supernatants were collected and protein concentration determined using the Bradford assay (BioRad Protein Assay Kit, USA). Proteins were homogenized in Sample buffer 1X in PBS [Laemmli, 5%(v/v)2-b-mercaptoethanol and 5% (v/v) bromophenol blue], boiled at 95°C for 5 minutes for protein denaturation and spinning at 14000rpm to clear the lysate.

40 µg of proteins were separated using sodium dodecyl sulphate-polyacrylamide gel electrophoresis (SDS-PAGE) on a 10% acrylamide gel (BIO-RAD) and transferred to nitrocellulose membrane by semi-dry blotting using the Trans-Blot Semi-Dry Transfer Cell (Bio-Rad) at 100 V for 2 h. Blocking with PBS containing 5% non-fat dry milk and 0.1 % Tween-20 was performed for 1 h at room temperature with agitation. Membranes were incubated with primary antibodies (Table 1) diluted in PBS-Tween-20 (PBS-T) with 1% non-fat dry milk overnight at 4 °C with agitation. After primary antibody incubation membranes were washed three times in PBS-T, for 15 min each, and incubated with an appropriate peroxidase-conjugated secondary antibody diluted 1:2000 in PBS-T for 1 h at room temperature with agitation. Membranes were then washed in PBS-T and incubated with 1 mL of Enhanced ChemiLuminescence (ECL) (Amersham) reagent for 5 minutes to allow protein visualization on an x-ray Amersham Hyperfilm ECL (GE Healthcare). Fiji-Image J Software was used for quantification of the difference in protein expression comparing with GAPDH or HSP70 expression.

**Table 1:** Western Blot primary antibodies.

| Primary antibody | Specie | Dilution | Seller | Reference |
| --- | --- | --- | --- | --- |
| Cleaved Caspase-3 (D3E9) | Rabbit | 1/1000 | Cell Signaling | 9579S |
| P-Cadherin (clone 56) | Mouse | 1/500 | BD Transduction | 610228 |
| E-Cadherin | Rabbit | 1/1000 | Cell Signaling | 24E10 |
| Vimentin (21H3) | Rabbit | 1/1000 | Cell Signaling | 5741T |
| Snail/Slug | Rabbit | 1/400 | Abcam | Ab85931 |
| Zeb2 | Rabbit | 0.1ug/ml | Invitrogen | PA5-82094 |
| GAPDH (0411) | Mouse | 1/1000 | Santa Cruz | Sc47724 |
| HSP70 | Mouse | 1/8000 | Santa Cruz | B-6 |

For cleaved-caspase-3 protein visualization, cells’ conditioned media was also collected together with cellular extracts, centrifuged at 2000rpm for 10min, in order to collect the dead cells and the cleaved caspase-3 fraction present in the conditioned media. Then protocol was carried out as explained above.

### siRNA transfection

P-Cadherin gene silencing (*CDH3*) was performed in MCF10A-Plk4p53KO using a validated siRNA, specific for *CDH3* (50nM, Hs_GCDH3_6), with the following target sequence 5’ AAGCCTCTTACCTGCCGTAAA 3′, from Qiagen (USA). P53 gene silencing was performed in MCF10A-Plk4, MCF10A-Plk4^1-608^ and RPE-Plk4 cell lines using a specific siRNA (100nM, L-003329-00-0020, Dharmacon). Transfections were carried out using Lipofectamine 2000 (Invitrogen), according to manufacturer’s recommended procedures. A scrambled siRNA targeting sequence 5’ AAGCCTCTTACCTGCCGTAAA 3′, with no homology to any gene, was used as a negative control (Qiagen, USA). Cells were incubated with the transfection mix for 6h. After siRNA transfection, cells were incubated for 24h in the presence or absence of Dox for 24h (1µg/ml) to induce extra centrosomes. Gene inhibition was evaluated by western blot after 31h of cell transfection for the MFE assay and after 47h for condition media collection.

### Condition media collection

For the conditioned medium assays, 5×10^5^ cells/well of MCF10A-Plk4 and MCF10A-Plk4^P53KO^ were grown until 80% - 85% confluence in 6 well plates. 24h after presence or absence of 1µg/ml Dox treatment for Plk4 overexpression, in serum-free media, conditioned media was collected, centrifuged at 1200 rpm for 5 min and filtered through a 0.2 µm pore filter. Condition media was then added to BT20 and MCF10A-Plk4^P53KO^ cells for 48 h.

### P-Cadherin ectodomain cleaved (soluble P-cadherin, sP-cad)

For P-Cadherin ectodomain cleaved detection, 5×10^5^ cells were grown until confluence in 0.2 mg/ml collagen-type I-coated 6 well plates (Merck, Darmstadt, Germany) and incubated in serum-free medium for 24 h with/without Dox treatment. For better detection of the soluble P-cadherin, serum-free conditioned media was filtered with 0.2*μ*m filter (Cytiva, Germany) for 5min at 1200rpm and concentrated using 30K Microcon Centrifugal Filter Devices (Merck, Darmstadt, Germany), according to manufacturer’s instruction. The proteins secreted were quantified in the recovered supernatant using the Bradford assay, and Western Blot was performed.

### qRT-PCR

qRT-PCR was performed in order to determine *PLK4* and P-Cadherin (*CDH3*) RNA expression. Cells were harvested and RNA extraction was performed using the RNeasy Kit (Qiagen, USA) according to manufacturer’s instructions. Concentration was determined in a ND-1000 spectrometer (Nanodrop) and 1 µg of total RNA was converted to cDNA using the Omniscript Reverse Transcriptase RT Kit (Invitrogen, USA). Quantitative Real-Time PCR (qRT-PCR) reaction was performed with TaqMan Gene Expression Assays (Applied Biosystems, USA), using gene-specific *PLK*4 and P-Cadherin (*CDH3*) PrimeTime probes (Integrated DNA Technologies, Inc., USA), recognizing specifically the corresponding cDNA sequences, which were amplified for 40 cycles. Analysis was performed with the 7500 Fast Real Time PCR Systems Instrument and software (Applied Biosystems), following the manufacturer’s recommendations. PrimeTime probe for *GAPDH* was also used as a housekeeping gene, and relative gene expression was determined by normalization. Data was analyzed by the comparative 2(-ΔΔCT). All reactions were done in triplicate and the results presented as mean of the values from three or more independent experiments. For *PLK4* mRNA detection on the MCF10A-Plk4, MCF10A-Plk4^p53KO^ and RPE-Plk4, a probe annealing on the exons 15-16 was used, while on MCF10A-Plk4^1-608^ Plk4 probe annealing on exons 1-3 was used, due to the truncation on the C-terminus (Table 2).

**Table 2:** List of probes used for qRT-PCR.

| Gene | Assay ID | Exon | RefSeq Number |
| --- | --- | --- | --- |
| *PLK4* | Hs.PT.58.2404526 | 15-16 | NM_001190799 |
| *PLK4* | Hs.PT.58.19849206 | 1-3 | NM_001190799 |
| *CDH3* | Hs.PT.58.2656432 | 7-8 | NM_001793 |
| *GAPDH* | Hs.PT.39a.22214836 | 2-3 | NM_002046 |

### Clonogenic assay

MCF10A-Plk4^p53KO^ cells were treated and not treated (negative control) with 1μg/ml of Doxycycline for 24h in order to induce Plk4 overexpression, and then harvested and replated equally (300 cells) in petri dish plates (7cm) and allowed to grow for 10days, where media was changed every 3days. Colonies were fixed with cold methanol and stained with 0,1% Sulforhodamine B (SRB), washed with PBS and air-dried followed by digital photography (Black-white image for better contrast visualization).

### Cell Viability

Cell viability was assessed using the PrestoBlue™ Cell Viability Reagent (Invitrogen, #A13262). One day prior to doxycycline induction, 2×10^4^ cells were seeded into a 96 multiwell plate. 24h after Plk4 overexpression, cells were washed twice with PBS and 50μL of 1:20 PrestoBlue Reagent was added (diluted in media). The plate was incubated for 30min at 37°C and fluorescence was read at excitation 569nm, Emission 590nm and 75% sensibility, using the BioTek’s SynergyTM MX microplate reader.

### Analysis of survival curves

For survival curves, Univariate survival curves were estimated with Kaplan-Meier and compared using the log-rank test. *P* values <0.05 were considered statistically significant.

## Acknowledgements

We thank Susana Godinho for the MCF10A-Plk4 and MCF10A-Plk4^1-608^ cell lines.

We also thank all the members of the Cancer Metastasis group and Cell Cycle Regulation (CCR) group (specially Sascha Werner, Mariana Faria and Paulo Duarte) for helping in the discussion of the work and for critical reading of the manuscript.

## Author contributions

I.F. carried out experiments, analyzed data and wrote the manuscript (original draft, review and editing). G.M. supervised the establishment and validation of the stable p53 knock-out cell line experiments. C.H. performed the epithelial and mesenchymal markers western blot and Plk4/P-cadherin qRT-PCR experiments, analyzed data and helped in discussion of the work. A.S.R. performed the analysis of Plk4 and CDH3 association in breast cancer cohort in the Kaplan Meier plotter. B.S. helped in the MFE and qRT-PCR analysis and discussion of the work. M.B.D. and J.P. supervised experiments, obtained funding and revised the manuscript. All authors read, revised and approved the final paper.

## Ethics Statement

This study did not require ethical approval.

