## supplementary files figures for "Polo-like kinase 4 (Plk4) potentiates *anoikis*-resistance of p53KO mammary epithelial cells by inducing a hybrid EMT phenotype"

**Supplementary Figure Legends**

**Supplementary Figure 1: p53 knock-out in the MCF10A-PLK4 cell line.** Western blot protein quantification of **(A)** p53 and its downstream target **(B)** p21 protein on MCF10A-Plk4 and MCF10A-PLK4^p53KO^ cells after exposure to 1 and 3 µM of Doxorubicin (DoxoR) in order to induce DNA damage (mild and severe DNA damage, respectively), for 4 hours, and 24h in drug free medium and 1 µg/ml of Doxycycline (Dox) for 24h. Integrated intensity was measured by ImageJ Software (National Institutes of Health). HSP70 was used as loading control, and all conditions were normalized to the control condition (MCF10A-Plk4 control). Data shown is average ± SD for three independent experiments, p<0.05, two-way ANOVA test.

**Supplementary Figure 2: Plk4 overexpression significantly potentiates cell viability of MCF10A-Plk4 and MCF10A-Plk4^P53KO^ cells.** Plk4 overexpression induced by Dox treatment (1μg/ml for 24h) induced a significand 0.5fold increase in the MCF10A-Plk4 and a 2.5fold increase in the MCF10A-Plk4^P53KO^ cell viability. This increase is more evident in the p53 knock-out background, demonstrating that the absence of p53 potentiates Plk4’s effect in cell viability. Data shown is average ± SD for four independent experiments, p<0.05, One-way ANOVA test.

**Supplementary Figure 3: Plk4 overexpression potentiates anoikis resistance of RPE-Plk4 cells in the absence of p53. (A)** *PLK4* mRNA expression levels in the RPE-Plk4 cell line after induction of Plk4 overexpression with Doxycycline treatment (1μg/ml of Dox for 24h). *GAPDH* mRNA expression was used as a housekeeping gene and condition was normalized to control (-Dox) condition. Data represents average ±SD for three independent experiments, p<0.05, Unpaired t test. (**B)** Quantification of centriole number in the RPE-Plk4 mitotic cells 24h of Dox treatment. Plk4 overexpression induced centriole amplification in 90% of cells 24h after Dox. To obtain the percentage of >4 centrioles, ≥100 cells were quantified per condition and per experiment. Data represents average ±SD for three independent experiments. p<0.05, Unpaired t test. **C)** *In vitro* quantification of Mammosphere Forming Efficiency (MFE) of RPE-Plk4 cells with and without Plk4 overexpression in the presence (siControl) and absence of p53 (sip53). The data is reported as the fold change in the percentage of mammospheres formed/7500 seeded cells ±SD, p<0.05, Two-Way ANOVA.

**Supplementary Figure 4: Conditioned medium (CM) from Plk4 overexpressed cells significantly increases *anoikis* resistance of MCF10A-Plk4^p53KO^ treated cells.** To promote Plk4 overexpression, MCF10A-Plk4^p53KO^ “CM donor” were treated with 1μg/ml of Dox for 24h, and MCF10A-Plk4^p53KO^ “CM receiver” cells were treated for 48 with CM from donor cells, following MFE assay. CM+Dox induced a significant increase in the MFE of MCF10A-Plk4^p53KO^ “CM receiver” cells when compared to control conditions (not treated and serum free media). Moreover CM-Dox did not induce an increase in MFE, suggesting that this effect is Plk4 mediated. Data represents three independent experiments and reported as the fold change in the percentage of mammospheres formed/7500 seeded cells ±SD, p<0.05, Two-WAY ANOVA.

**Supplementary Figure 5: Co-expression** **of high Plk4 and high P-cadherin expression in breast tumors are significantly associated with worse prognosis.**

Patients with breast tumors co-expressing high Plk4 and P-cadherin expression showed a significant correlation with worse DFS and OS, as revealed in Kaplan-Meier plots. Univariate survival curves were estimated with Kaplan-Meier and compared using the log-rank test. *P* values <0.05 were considered statistically significant**.**
