## Supplementary figures and images for "Polo-like kinase 4 (Plk4) potentiates *anoikis*-resistance of p53KO mammary epithelial cells by inducing a hybrid EMT phenotype"

### supplementary file 1

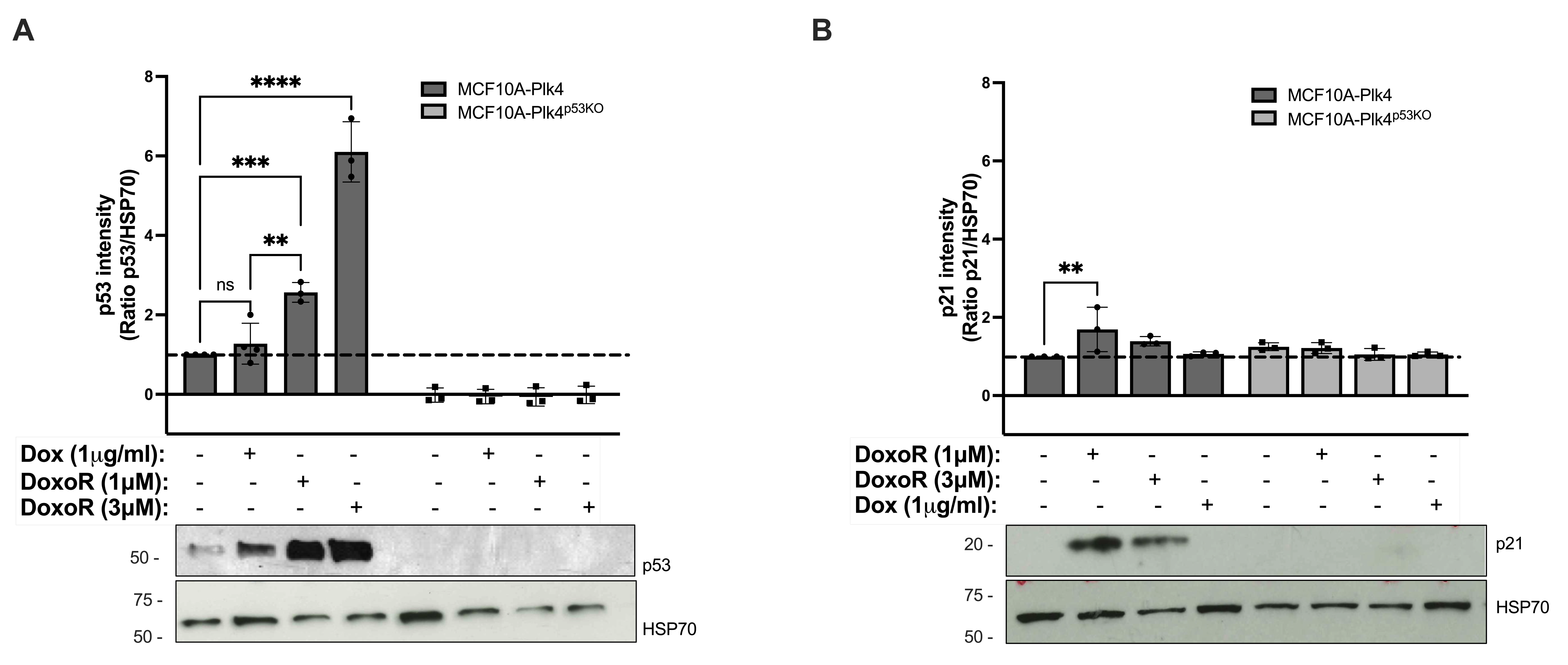

### supplementary file 2

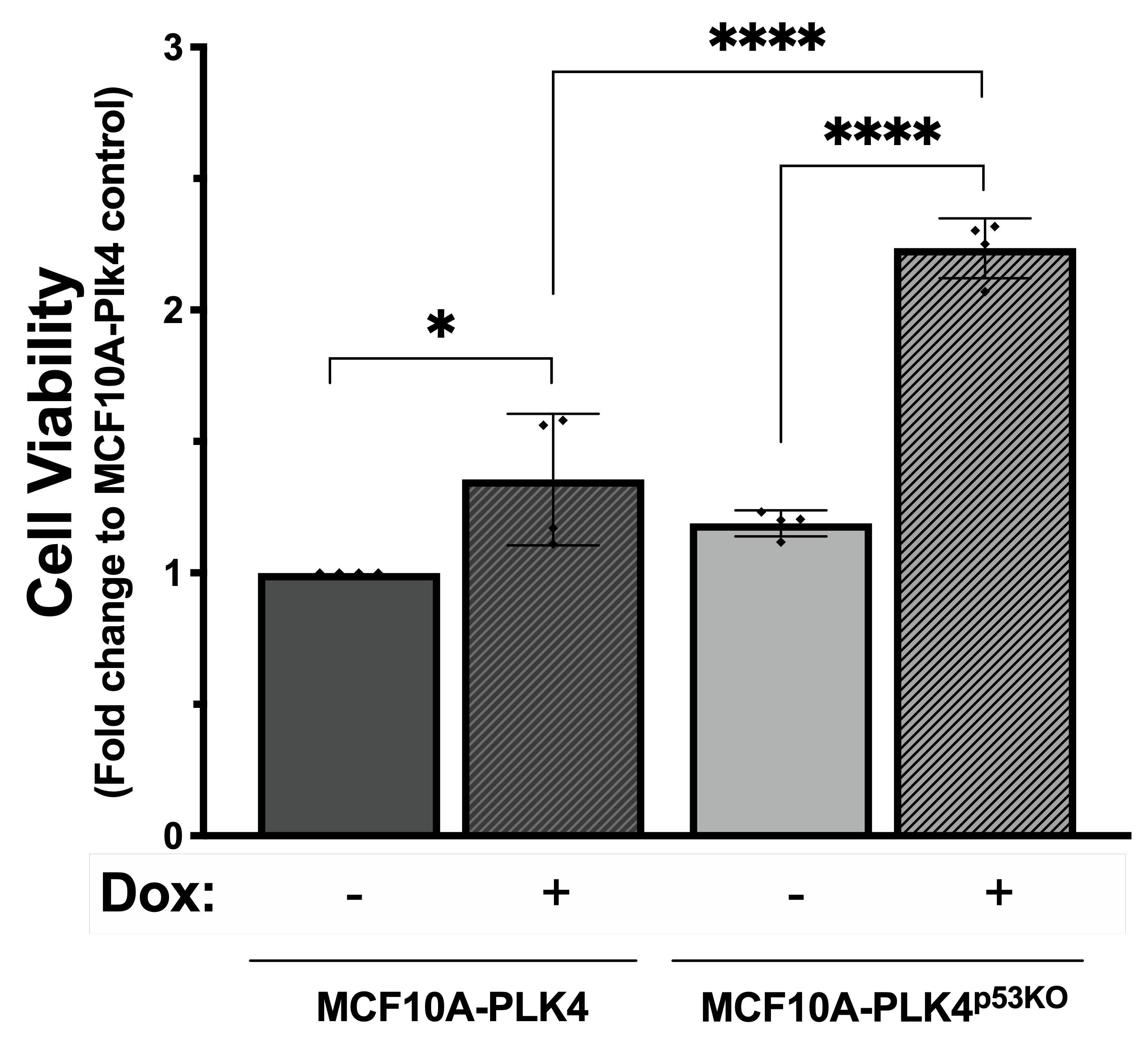

### supplementary file 3

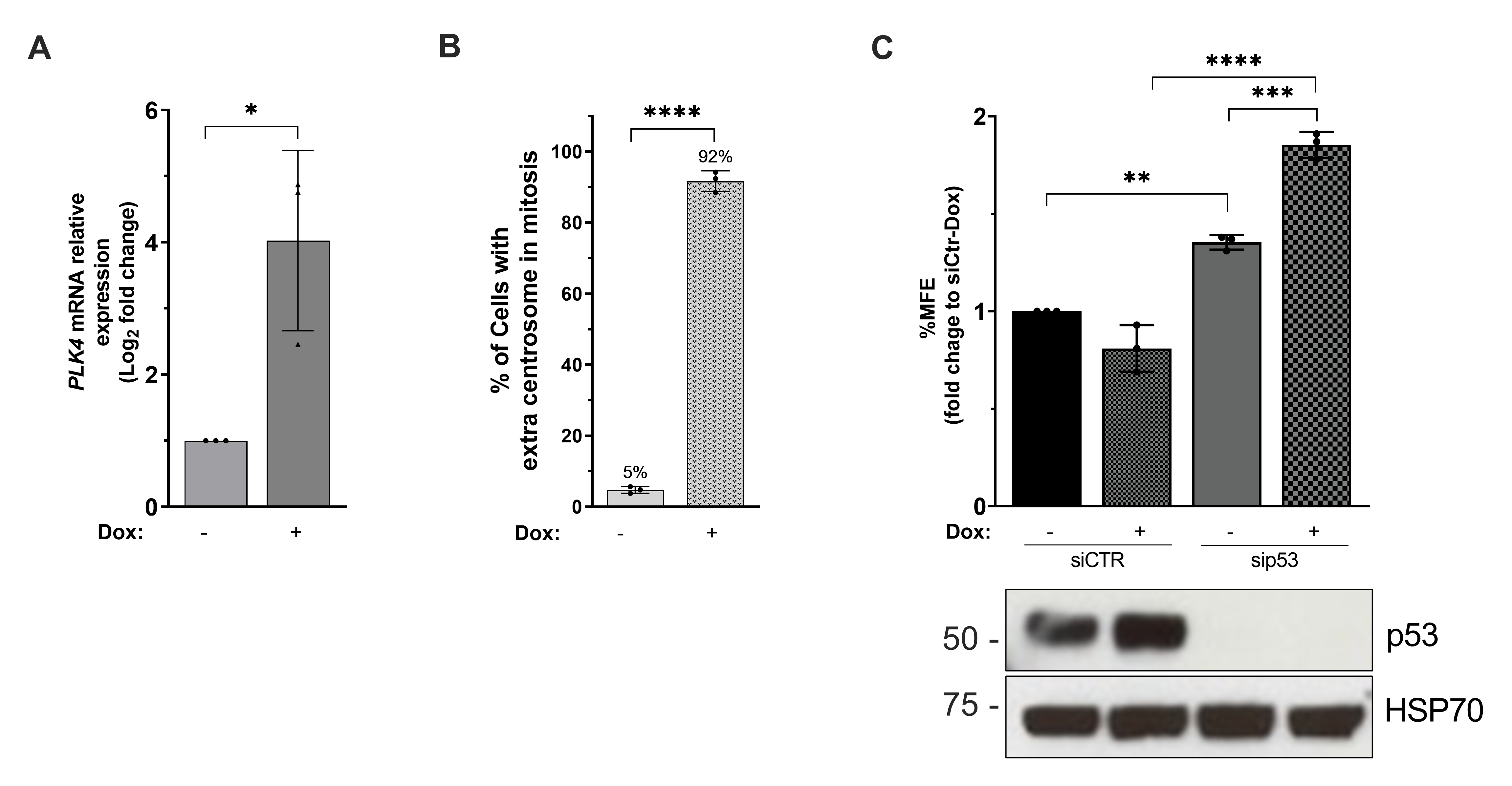

### supplementary file 4

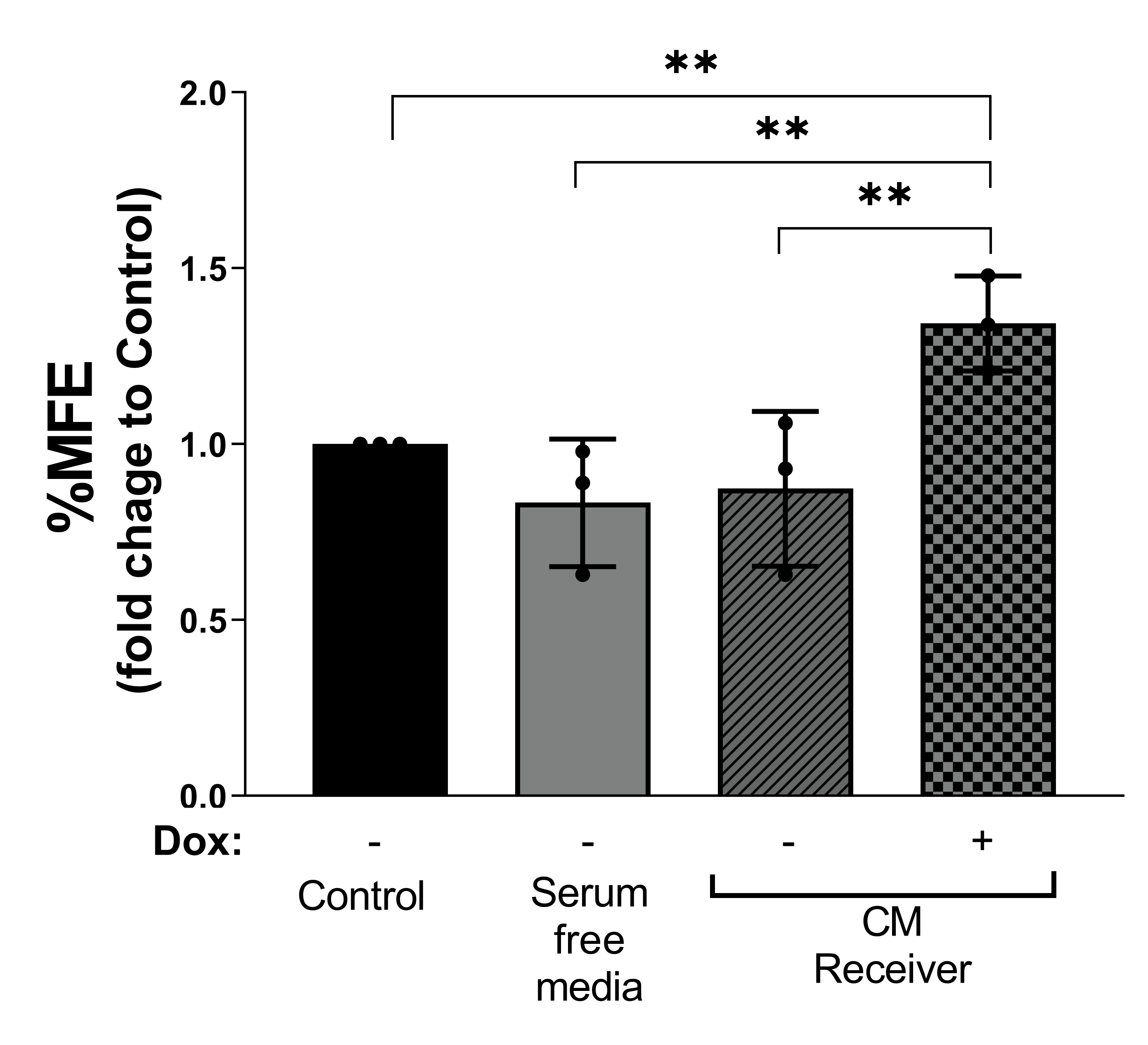

### supplementary file 5

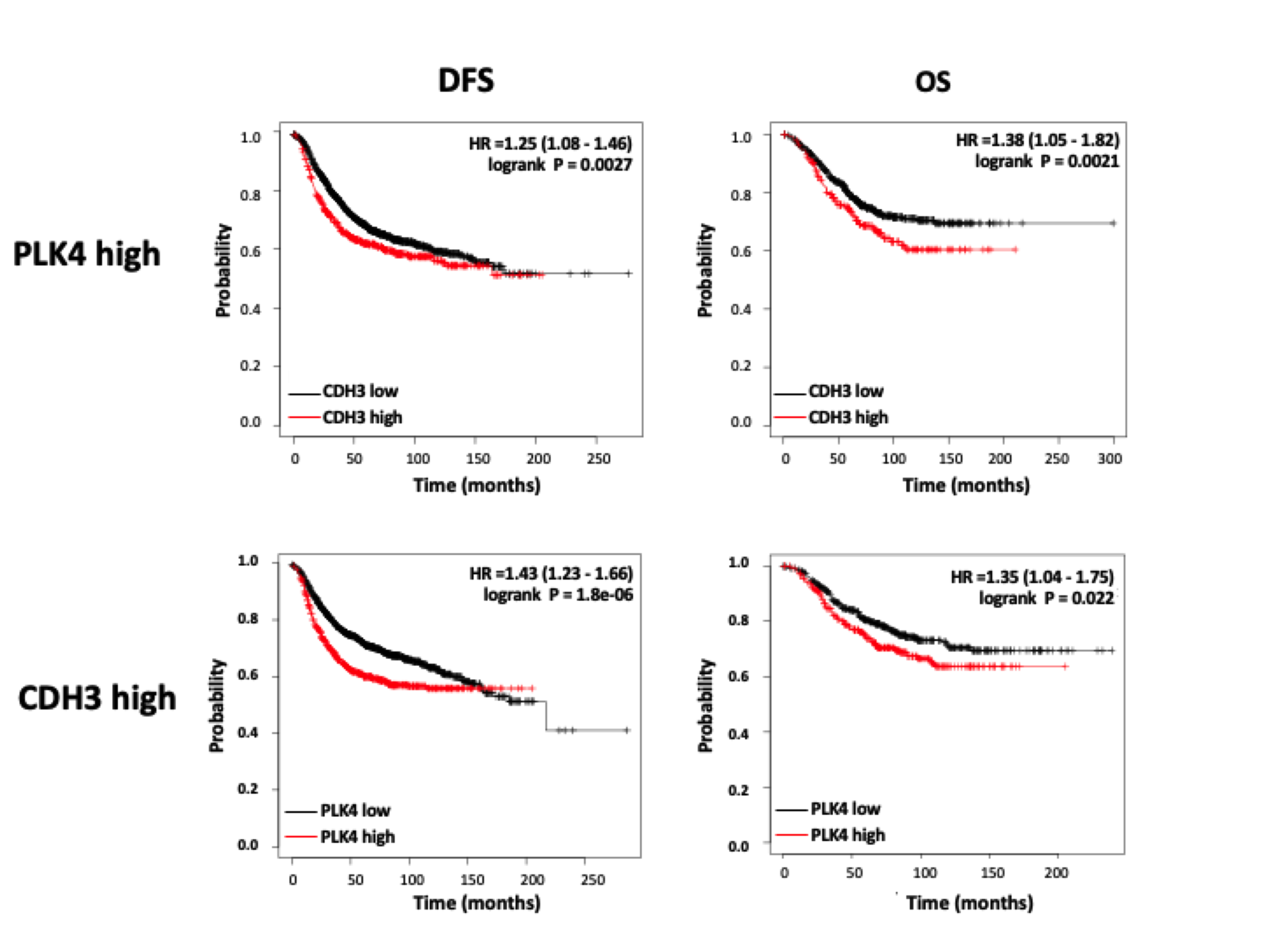
